# L-DOPA dioxygenase of the fly agaric toadstool: revision of the *dodA* gene sequence and mechanism of enzymatic pigment production

**DOI:** 10.1101/2020.08.03.235077

**Authors:** Douglas M. M. Soares, Letícia C. P. Gonçalves, Caroline O. Machado, Larissa Cerrato Esteves, Cassius V. Stevani, Carla C. Oliveira, Felipe A. Dörr, Ernani Pinto, Flávia M. M. Adachi, Carlos T. Hotta, Erick L. Bastos

## Abstract

l-DOPA extradiol dioxygenases (DODAs) catalyze the production of betalains and hygroaurins pigments. The sequence of the DODAs found in Caryophyllales and Basidiomycetes are not conserved, although betalains are produced both by plants and fungi. Here we revise the coding region of the *dodA* gene of fly agaric [*Amanita muscaria* (L.) Lam.] and describe an alternative start codon downstream that enables the heterologous expression of AmDODA, a promiscuous l-DOPA dioxygenase. AmDODA is 43-amino acid residues shorter than the recombinant DODA previously reported but catalyzes the formation of two isomeric seco-DOPAs that are the biosynthetic precursors of betalains and hygroaurins. The putative active site of AmDODA contains two distinct His-His-Glu motifs that can explain the dual cleavage of l-DOPA according to the mechanism proposed for non-heme iron-dependent dioxygenases. Upon addition of excess l-DOPA, both the betaxanthin and hygroaurin adducts of l-DOPA are produced. The kinetic parameters of enzymatic catalysis at pH 8.5 are similar to those reported for other l-DOPA dioxygenases. The rate constants for the conversion of l-DOPA into the betalamic acid and muscaflavin were estimated by kinetic modelling allowing the proposal of a mechanism of pigment formation. These results contribute to understanding the biosynthesis of bacterial, fungal and plant pigments, for the biotechnological production of hygroaurins, and for the development of more promiscuous dioxygenases for environmental remediation.

## 1. INTRODUCTION

The biological and ecological functions of pigments in plants and fungi are more complex than meets the eye (Davies *et al*., 2018). Most Caryophyllales plants and a few Basidiomycete fungi of genera *Amanita, Hygrocybe* and *Hygrophorus* produce betalains (Lodge *et al*., 2013; Polturak & Aharoni, 2018), which impart to these organisms their bright red-violet and orange colors and, occasionally, green fluorescence (Gandía-Herrero *et al*., 2005). Fungi produce neither anthocyanins nor any other flavonoid (Gil-Ramírez *et al*., 2016), and the occurrence of betalains and anthocyanins in plants would be considered redundant since their colors and functions against biotic and abiotic stress are similar (Davies, 2015; Osbourn, 2017). Interestingly, betalains and anthocyanins were never found in the same organism in the wild (Stafford, 1994; Brockington *et al*., 2011).

Adding to the phylogenetic importance of these observations, the biosynthesis of betalains in plants and fungi depend on enzymes that have sequences with no identities and, thus, different evolutive origins (Christinet *et al*., 2004). In several living organisms, l-tyrosine is hydroxylated in the presence of cytochrome P450-like enzymes producing 3,4-dihydroxy-l-phenylalanine (l-DOPA) (Hatlestad *et al*., 2012; Polturak *et al*., 2016; Wei *et al*., 2018). In plants, the oxidative cleavage of the catechol ring of l-DOPA is catalyzed by non-heme 4,5-extradiol dioxygenases, ultimately leading to betalamic acid, which is the biosynthetic precursor of betalains (Fig. 1) (Timoneda *et al*., 2019). The fungal enzyme catalyzes the oxidative cleavage of the catechol ring at both positions 4,5 and 2,3 (Strack *et al*., 2003; Vaillancourt *et al*., 2006). The resulting seco-DOPA derivatives, viz., muconate 6-semialdehydes or α-pyrone amino acids, are highly functionalized species that contain several reactive functional groups. Consequently, the product of cleavage of l-DOPA at the 4,5 position, betalamic acid, and its two constitutional isomers originated from the 2,3-breakage can be produced (Fig. 1). Muscaflavin is the precursor of the poorly understood and rare hygroaurins. l-4-(2-oxo-3-butenoic-acid)-4,5-dihydropyrrole-2-carboxylic-acid (OBDC) (Barth *et al*., 1979; Saha *et al*., 2015), on the other hand, is the key intermediate for the biosynthesis of antitumor pyrrolo[1,4]benzodiazepines, the bacterial hormone hormaomycin, and the lincosamide antibiotic lincomycin (Jiraskova *et al*., 2016). Betalamic acid, muscaflavin and OBDC show absorption maxima at 414 nm at pH 8 (Saha *et al*., 2015). Last, the combined action of an NADP^+^- and Zn^2+^-dependent l-DOPA 2,3-dioxygenase and stizolobinate synthase produce stizolobinic and stizolobic acids in the velvet bean [*Mucuna pruriens* (L.) DC.] from l-2,3- and l-4,5-seco-DOPA, respectively (Saito & Komamine, 1978; Musso, 1979).

**Fig. 1.**
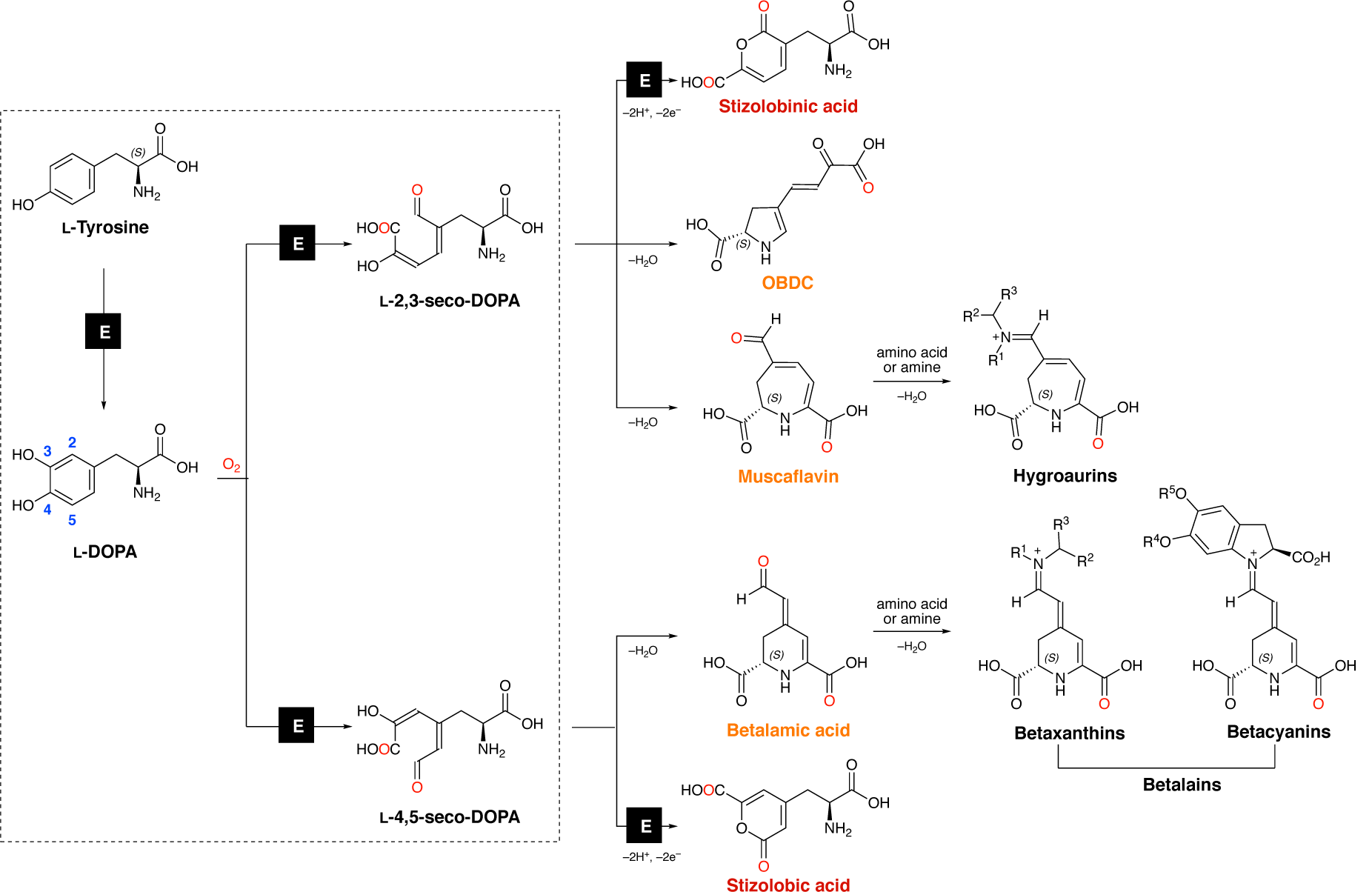
Biosynthesis of betalains and hygroaurins. The formation of the two classes of betalains, *viz*., betacyanins and betaxanthins, both *in vivo* and *in vitro* are the result of the spontaneous coupling between betalamic acid and either cyclo-DOPA derivatives or amines and amino acids, respectively. The analogous reaction of amines or amino acids with muscaflavin produces hygroaurins. The dehydrative cyclization of l-4,5-seco-DOPA is spontaneous in acidic aqueous conditions (Schliemann *et al*., 1999). Color code indicates isomers and the symbol “E” designates enzymatic transformations. Amino and carboxyl groups are presented in the uncharged forms for clarity.

The *dodA* gene (GenBank Y12886, 1629-bp) encodes the DOPA extradiol dioxygenase (DODA_AMAMU: Uniprot P87064) of the toadstool fly agaric [*Amanita muscaria* (L.) Lam.]. After the description of the *dodA* gene by Zrÿd and coauthors, the genetic basis of betalain biosynthesis started to unravel (Hinz *et al*., 1997). The same group screened a cDNA library from the red-colored pileus (cap tissue) of *A. muscaria* by using anti-DOPA dioxygenase antibodies and found twenty positive clones, all having a 612-bp open reading frame (ORF) and a truncated 5’ end (Mueller *et al*., 1997b). Although the first 27-aa residues were not encoded by the cDNA clones, the resulting recombinant 228-aa dioxygenase was active to produce both betalamic acid and muscaflavin, and show similar activity compared to the native enzyme (Mueller *et al*., 1997a; Mueller *et al*., 1997b). Despite the obvious economic advantages of producing natural and pseudo-natural betalains (Freitas-Dörr *et al*., 2020), hygroaurins and OBDC derivatives, their chemoenzymatic synthesis using the recombinant *A. muscaria* dioxygenase was developed no further.

We amplified and cloned the coding sequence (CDS) of the *dodA* gene from the pileus of *A. muscaria*. DNA sequencing showed that the 3’ canonical splice site of the first intron was not AG, as expected (Hinz *et al*., 1997), but GA. The occurrence of GA in this position causes the retention of the first intron, leading to a truncated protein of 35-aa residues. Consequently, finding an alternative start codon is required for the enzyme expression.

Here we report the cloning of the CDS of the *A. muscaria*’s l-DOPA dioxygenase from an alternative start codon and the expression of a 205-aa recombinant enzyme AmDODA that is active to produce both betalamic acid and muscaflavin from l-DOPA. The ATG start codon is located at the second exon of the *dodA* gene, downstream to the previously annotated one. Ascorbic acid was used to prevent the non-enzymatic oxidation of l-DOPA. The kinetics of betalain and hygroaurin formation from l-DOPA was investigated by HPLC-DAD-ESI-QTOF-MS/MS and the rate constants for the elementary steps in the synthesis of betalamic acid and muscaflavin were estimated by kinetic modelling. These new findings allow the heterologous expression of AmDODA, contribute for the overall understanding of the biosynthesis of high valued secondary metabolites, enable the biotechnological production of muscaflavin, and facilitate the metabolic engineering of plants able to produce fungal pigments.

## 2. MATERIALS AND METHODS

### 2.1. Molecular biology

#### 2.1.1. Cloning of the coding sequence for *dodA* gene

A*manita muscaria* mushrooms were collected in Santana de Parnaíba, São Paulo, Brazil (23°28’18.8’’ S 46°51’50.2’’ W) on June 20, 2018 (Fig. 2a). DNA and mRNA samples were extracted from the red-pigmented pileus using the DNeasy® and RNeasy® kits (QIAgen). cDNA was synthesized from mRNA (1 µg) by using the SuperScript™ III reverse transcriptase (ThermoFisher Scientific). Primer design was carried out using the DNA sequence of the *A. muscaria dodA* gene (GenBank Y12886) (Hinz *et al*., 1997). The coding sequence of *dodA* was PCR amplified from DNA and cDNA samples using the primers dodA-F (CACCATGGTGCCAAGCTTCGTTGT) and dodA-R (CTATGCATCTCGATGGGGCGCTCT). PCR was carried out under standard conditions using the Phusion High-Fidelity DNA Polymerase (ThermoFisher Scientific). Samples were kept at 98 °C for 30 s (1 cycle), 98 °C for 10 s (30 cycles), 62 °C for 30 s (1 cycle), and 72 °C for 1 min (1 cycle), followed by a final extension phase at 72 °C for 7 min. Products were cloned into pENTR™/SD/D-TOPO™ (ThermoFisher Scientific) and transformed in DH5α competent *E. coli* (ThermoFisher Scientific), according to the instructions of the manufacturer. The sequence was checked by DNA sequencing of the positive colonies using the primers M13 forward (5’-GTAAAACGACGGCCAG-3’) and M13 reverse (5’-CAGGAAACAGCTATGAC-3’). Due to the results obtained, a new coding sequence was proposed for *dodA* gene, hereby named *AmDODA*, which was deposited in the NCBI database under the GenBank accession number MK922469. cDNA samples from *A. muscaria* were used for PCR amplification of the *AmDODA* using the AmDODA-F (5’-ACTTTAAGAAGGAGATATACATGTCCACCAAGCCAGAG-3’) and AmDODA-R (5’-GTCGACGGAGCTCGAATTCGGTGCATCTCGATGGGGCG-3’) primers. PCR was carried out under standard conditions using the Q5^®^ High-Fidelity DNA Polymerase (New England Biolabs). Samples were kept at 98 °C for 30 s (1 cycle), 98 °C for 10 s (30 cycles), 60 °C for 30 s (1 cycle), and 72 °C for 1 min (1 cycle), followed by a final extension phase at 72 °C for 2 min. PCR products were cloned in the pET28b vector (Novagen) linearized with *Bam*HI and *Not*I according to the sequence and ligation-independent cloning (SLIC) method (Jeong *et al*., 2012). The recombinant plasmid pET28b-*AmDODA* was confirmed by DNA sequencing using the primers T7 promoter (5’-TAATACGACTCACTATAGGG-3’) and T7 terminator (5’-GCTAGTTATTGCTCAGCGG-3’).

**Fig. 2.**
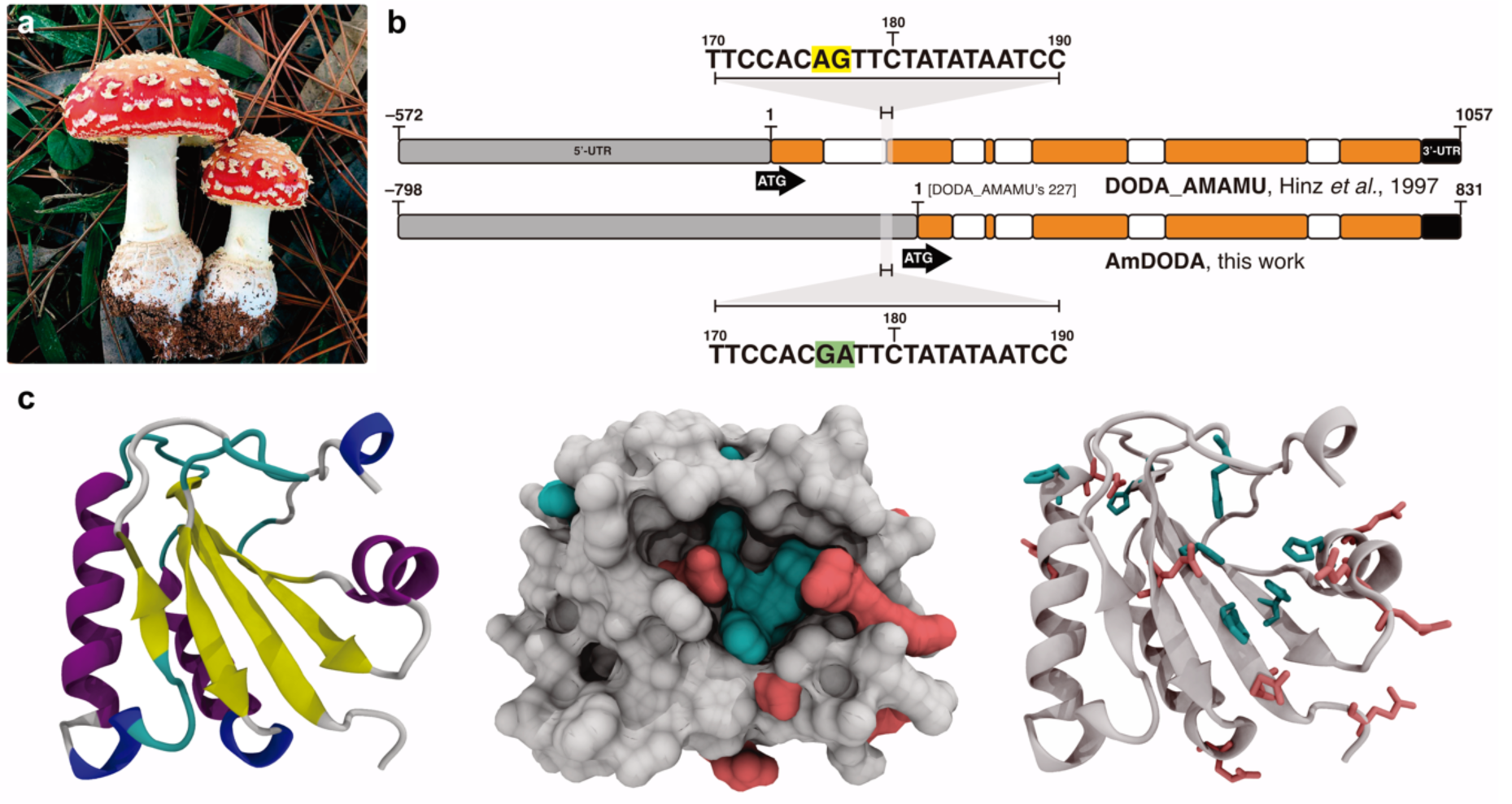
Fly agaric, CDS of the *AmDODA* gene and *in silico* structure of AmDODA. (a) Fly agaric (*A. muscaria*) used for the extraction of genetic material, which was found in pines trees (*Pinus* sp. L.). (b) Comparison of the nucleotide sequence of *dodA* and *AmDODA* genes that codify for the DODA_AMAMU and AmDODA proteins, respectively. The expanded region highlights the modification in the canonical 3’ splice site. The start codon is indicated by the ATG sequence. Blocks in orange are exons. The complete sequence of *dodA* is presented in the *Supporting Information*. (c) *In silico* model of the DODA_AMAMU and AmDODA proteins obtained using the Phyre^2^ server. Representation of secondary structures and solvent accessible surface (140 pm); GLU and HIS are highlighted in pink and cyan, respectively.

#### 2.1.2. Expression and purification of AmDODA

*Escherichia coli* strain BL21 (DE3) (New England Biolabs) chemically competent was transformed by heat-shock with the recombinant plasmid pET28b-*AmDODA* to express the C-terminal His-tagged AmDODA. 2-YT medium (3 mL) supplemented with kanamycin (50 μg mL^−1^) was inoculated with *E. coli* BL21(DE3) pET28b-*AmDODA* and grown overnight at 37 °C in an orbital shaker operated at 200 rpm. The cultures were transferred to 1.0 L baffled Erlenmeyer flask containing 250 mL 2-YT/kanamycin medium and shaken at 200 rpm and 37 °C for approximately 2 h to a final optical density of 0.4 to 0.6 at 600 nm. Isopropyl β-D-thiogalactopyranoside (IPTG) was added to a final concentration of 0.5 mM for the induction of AmDODA expression and the flasks were incubated for 16 h at 30 °C. Cells were harvested by centrifugation (8000 ×g, 4 °C, 30 min) and resuspended in sodium phosphate lysis buffer (Supporting Information). Cell lysis was performed in a French Press Cell G-M™ (ThermoFisher Scientific) and the recombinant protein was purified by gravity-flow chromatography using a nickel-charged resin Ni-NTA Agarose (QIAgen) equilibrated with 10 mM imidazole in the lysis buffer. An elution buffer (linear gradient of imidazole from 100 mM to 500 mM) was used to elute AmDODA. Fractions containing the enzyme were identified by sodium dodecyl sulfate polyacrylamide gel electrophoresis (SDS-PAGE) analysis (Laemmli, 1970), by application to 15% polyacrylamide gels and stained using a standard Coomassie blue method. Pure fractions were pooled and desalted by dialyzes in sodium phosphate buffer (50 mM, pH 7.4). Protein concentrations were determined by the dye-binding method of Bradford assay (Bio-Rad) and bovine serum albumin as the calibration standard. The revised nucleotide sequence was deposited in the GenBank database under the accession code MK922469 (Soares *et al*., 2019).

### 2.2. Enzyme catalysis

The oxidation of l-DOPA in the presence of AmDODA was monitored at every 18 s for 5 min by UV-Vis absorption spectroscopy (300 – 600 nm, scan rate: 2,400 nm min^−1^) using a Varian Cary 50 Bio spectrophotometer equipped with a cell holder thermostated at 25 °C. The reaction was initiated by adding the enzyme (1 μM) to a solution of l-DOPA (1 mM) and ascorbic acid (10 mM) in sodium phosphate buffer (50 mM, pH 8.5) unless otherwise stated. Since the addition of AscH decreases the medium pH, careful correction with base must be performed before the addition of the substrate. Product formation was monitored by the increase in absorption at 414 nm over time and the initial rate of product formation (in μM min^−1^) was calculated by linear regression assuming a molar absorptivity coefficient (ε) at 424 nm for both products of 24,000 M^−1^ cm^−1^ (Trezzini & Zrÿb, 1991; Contreras-Llano *et al*., 2019). The specific enzyme activity was then calculated by dividing the initial rate of product formation (in μM min^−1^) by the initial enzyme concentration (in mg L^−1^) and, thus, corresponding to the molar quantity of product converted each minute per mass of enzyme, i.e., μmol min^−1^ mg^−1^ or U mg^−1^ (Harris & Keshwani, 2009). The specific activity at each substrate concentration (l-DOPA, 0.5 to 7 mM range) was plotted and the values of *K*M and Vmax were calculated by the non-linear fitting of the data to the Michaelis-Menten equation, without considering substrate inhibition. All regressions were carried out using the Origin 2016 software (OriginLab).

### 2.3. Apoenzyme preparation

Sodium phosphate buffer (50 mM, pH 8.5) was treated with Chelex-100 (50 mg mL^−1^) for 24 h and the pH was confirmed and adjusted when necessary (Ch-PB). Stock solutions of ascorbic acid (250 mM) and l-DOPA (5 mM) were prepared using the Ch-PB. The stock solution of AmDODA in sodium phosphate buffer (1.12 mg mL^−1^, 50 μM, 10 μL) was diluted 100-fold using Ch-PB and Chelex-100 (50 mg) was added. The pH of the supernatant was checked before use. The mixture was incubated at 4 °C and 450 rpm using an orbital mixer, and aliquots were taken every 30 min until the activity of the enzyme, treated with Ch-PB, was lost (3.5 – 5 h). The activity of the enzyme prepared with Ch-PB was compared to that of the negative control; all solutions were prepared using sodium phosphate buffer and incubated at the same condition.

### 2.4. *In silico* enzyme modelling

The amino acid sequences of AmDODA and AMAMU-DODA (UniProt ID P87064) were submitted to structural homology modelling using the Phyre^2^ server (Kelley *et al*., 2015). For both proteins, the crystal structure of the putative dioxygenase YP_555069.1 from *Burkholderia xenovorans* strain LB400 (140 pm resolution, RCSB PDB: 2P8I) was selected as the highest score template for the initial *in silico* structural modeling (Rank: 1; Aligned residues: 109; Identity: 40%; Confidence: 100%). The profile-based threading method program Phyre^2^ was able to model 64% of the sequence with > 90% of confidence.

### 2.5. Chromatographic analysis of product formation

The reaction of l-DOPA (2.5 mM) and oxygen in the presence of ascorbic acid (10 mM) and AmDODA (1.0 μM) in sodium phosphate buffer (50 mM, pH 8.5) at 25 °C was monitored over time by HPLC-PDA using a Shimadzu Prominence liquid chromatograph equipped with an Ascentis C18 column (5 μm, 250 × 4.6 mm, Supelco) and a SPD-M20A detector. The reaction was analyzed at 1.0 mL min^−1^ and at 25 °C under (condition 1) a linear gradient from 2 to 60% B in 20 min (solvent A: water; solvent B: acetonitrile, both containing 0.05% v/v formic acid) or (condition 2) isocratic 5% B for 5 min then a linear gradient from 5% to 25% B in 15 min. After equilibrium was reached, the reaction mixture submitted to HPLC-HRMS analysis using a Shimadzu Prominence liquid chromatograph equipped with a Luna C18 column (3 μm, 150 × 2 mm, Phenomenex^®^) and coupled to a Bruker Daltonics microTOF-QII mass spectrometer fitted with an electrospray source operated in positive mode. The reaction mixture analyzed at 0.2 mL min^−1^ at 30 °C under a linear gradient from 5 to 95% B in 15 min (solvent A: 0.05% v/v formic acid in water, solvent B: 0.05% v/v formic acid in acetonitrile). l- and D-DOPA, betalamic acid and dopaxanthin were used as standards (*see the Supporting Information*). HBt was quantified by absorption spectroscopy using a molar absorption coefficient at 424 nm of 24,000 M^−1^ cm^−1^ (Contreras-Llano *et al*., 2019).

### 2.6. Kinetic modelling

To model the kinetics of 2,3-seco-DOPA, 4,5-seco-DOPA, betalamic acid, muscaflavin, l-dopaxanthin and l-DOPA hygroaurin formation, the observed rate constants (*k*_obs_) for the formation and/or consumption of these intermediates and products were calculated by the non-linear fitting of the chromatographic data to mono- or biexponential functions using the Origin 2016 software (OriginLab). Next, a minimal kinetic model was used to investigate the conversion of 1.0 mM l-DOPA into l-dopaxanthin and l-DOPA hygroaurin using COPASI v.4.22 (Hoops *et al*., 2006). For simplicity, all elementary reactions were defined as unimolecular and irreversible and the values of *k*_obs_ were used to develop the model ensuring that the resulting kinetic profile was qualitatively similar to the experimental data. The maximum theoretical concentration of each intermediate and product was divided by the maximum experimental absorption measured by HPLC-PDA analysis. The resulting factor was used to calculate the relative concentration of each species from its experimental absorption. Changes in the concentration of each species over time were fitted to different kinetic models and the first-order rate constants were calculated using the evolution programming method.

## 3. RESULTS AND DISCUSSION

### 3.1. Cloning, expression and purification of AmDODA

The CDS for the l-DOPA extradiol dioxygenase of *A. muscaria* in the *dodA* gene is 687 bp long (GenBank: Y12886.1, Fig. S1). However, our amplification of the *dodA* CDS resulted in PCR products of 784 bp. We inferred that this discrepancy could be explained by the occurrence of alternative splicing or by the use of a different variety of *A. muscaria* for the extraction of genetic material. The variety of *A. muscaria* used by Zrÿd and coauthors was collected in the Jorat forest, near Lausanne (Hinz *et al*., 1997) and, therefore, should correspond to the Euro-Asian fly agaric [*A. muscaria* var. *muscaria* (L.) Lam.]. The subspecies used in this work (Fig. 2a) is either the Euro-Asian or the American fly agaric (*Amanita muscaria* subsp. *flavivolvata* Singer) (Wartchow *et al*., 2013), both of which produce betalains and hygroaurins. DNA sequencing and alignment with the reported mRNA sequence for *dodA* revealed the retaining of the first intron, which is 97 nt long, viz., 784 nt – 687 nt. Furthermore, a GA was found at the 3’ splice site of the first intron, differing from the canonical AG reported previously (Hinz *et al*., 1997) (Fig. 2b). Intriguingly, the DNA fragment obtained from the *dodA* CDS deposited in the GenBank would result in a truncated polypeptide of 35-aa residues, if the AG was mutated to GA, causing an intron retention, impelling us to seek for alternative start codons.

All the 14 putative initiation ATG codons found in the genomic sequence lead to truncated polypeptides, except for the one located at the position 227 of the Exon 2, which remains in the same reading frame of the published *dodA* CDS (Fig. 2b). The ATG at position 227 is embedded in a close to consensus Kozak sequence, whereas the first ATG is not, which could indicate that translation would start more efficiently at the ATG 227. The *AmDODA* CDS starting from this alternative start codon is 558 bp long and codes for a 185-aa protein (Fig. S2). *E. coli* BL21 (DE3) cells transformed with the recombinant plasmid pET28b-*AmDODA* were incubated with 0.5 M IPTG at 30 °C overnight producing a recombinant 205-*aa* His-tagged dioxygenase that was active to produce both betalamic acid and muscaflavin from l-DOPA. This recombinant enzyme, hereafter called AmDODA, has the same primary sequence of the DODA_AMAMU dioxygenase reported by Zrÿd and coauthors (Hinz *et al*., 1997) except for the absence of the first 43-aa residues (Fig. 2b) that, coincidentally, are absent in other fungal dioxygenases (Fig. S3). Indeed, the absence of the first 30-aa was reported to not affect the overall stability and activity of the enzyme (Hinz *et al*., 1997; Mueller *et al*., 1997b).

*In silico* modelling of the structures of AmDODA and DODA_AMAMU structure converged to the same minimal tridimensional model, which is based on the putative dioxygenase of *Burkholderia xenovorans* strain LB400 (Chain *et al*., 2006) (Fig. 2c). The final model does not contain any metal cation in the active site, even though most catechol 2,3- and 4,5-extradiol dioxygenases depend on iron(II) as a cofactor to enable triplet molecular oxygen to react rapidly with singlet substrates (Bugg & Winfield, 1998; Bugg, 2003; Bugg & Ramaswamy, 2008; Wang *et al*., 2017). Nevertheless, several proximal His-His-Glu motifs, which are involved in Fe(II) complexation at the active site of several extradiol dioxygenases (Vaillancourt *et al*., 2006; Bugg & Ramaswamy, 2008), were found at the surroundings of a putative catalytic pocket (Fig. 2c).

### 3.2. Enzyme kinetics and synthesis of betalamic acid and muscaflavin

#### 3.2.1. Effect of cofactors

Several cofactors have been used to promote reactions involving l-DOPA dioxygenases (Girod & Zryd, 1991; Terradas & Wyler, 1991; Mueller *et al*., 1997b; Gandía-Herrero & García-Carmona, 2012; Grewal *et al*, 2018; Contreras-Llano *et al*., 2019). To constrain the complexity of the system, we investigate the catalytic activity of AmDODA only in the presence of ascorbic acid (AscH), which prevents the oxidation of the substrate by oxygen, thus increasing the overall efficiency of the biocatalytic process.

The addition of l-DOPA leads to an increase in the absorption at around 420 nm without producing browning substances that are formed in the absence of AscH (Figs. 3a, S4 and S5). AscH is an important cofactor in iron- and 2-oxoglutarate-dependent dioxygenases (Kuiper & Vissers, 2014), favoring the enzyme turnover by reducing enzyme-bound Fe(III) to Fe(II) formed during enzymatic activity and, most importantly, by uncoupled reaction cycles (Kuiper & Vissers, 2014). Ascorbate is also efficient enhancing the activity of oxygenases and dioxygenases compared to other reducing agents, including DTT and NADPH, due to its specific binding to the active site of the enzyme (Nietfeld & Kemp, 1981; De Jong *et al*., 1982; Wu *et al*., 2016). Nevertheless, the addition of AscH does not affect the initial activity of AmDODA (Fig. S4).

**Fig. 3.**
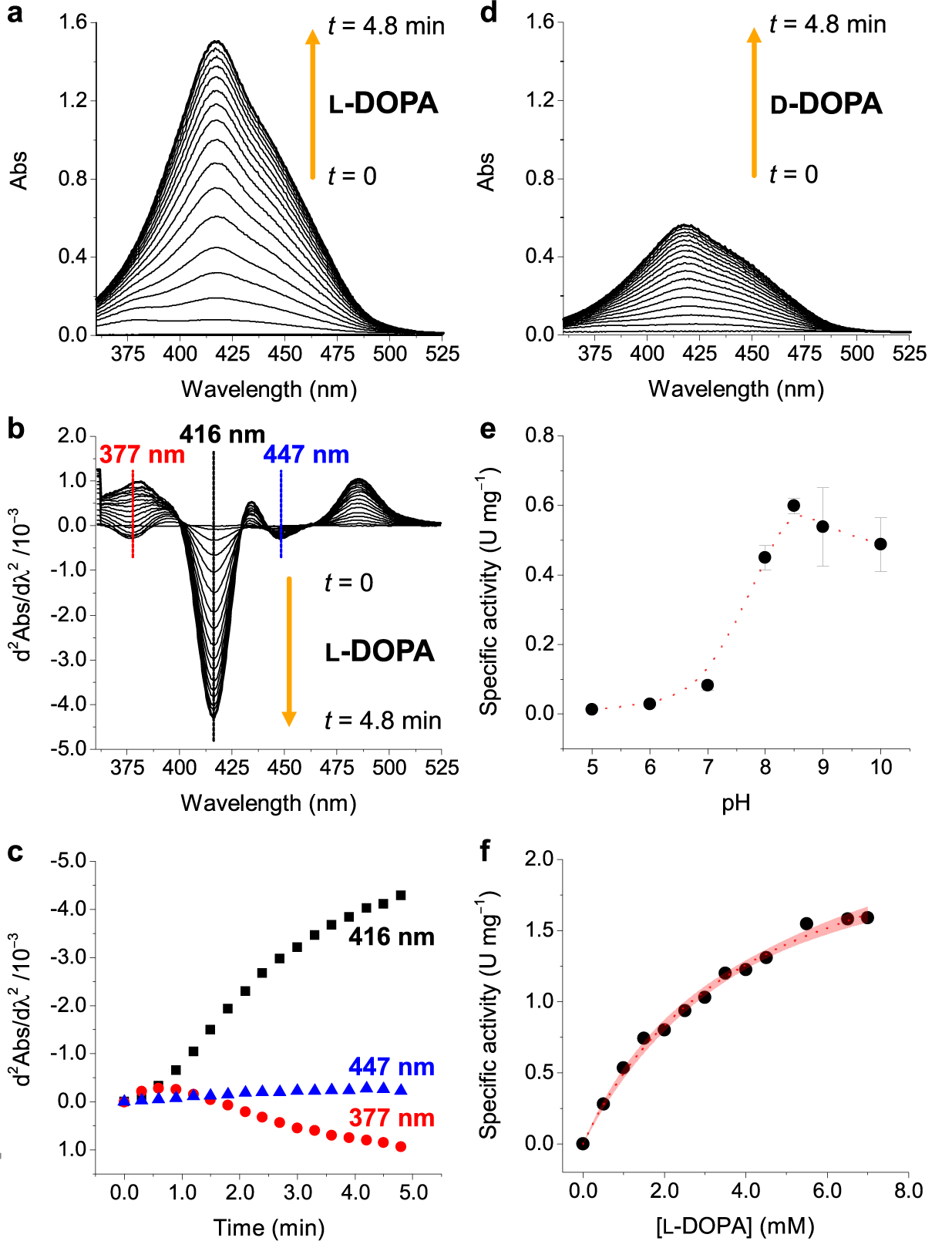
Enzyme activity studies. (a) UV-Vis spectra of the oxidation of l-DOPA by oxygen catalyzed by AmDODA over time. (b) Second derivative UV-Vis spectroscopy and (c) kinetic tracing at three representative wavelengths. (d) UV-Vis spectra of the oxidation of D-DOPA by oxygen catalyzed by AmDODA over time. (e) Effect of the pH on the AmDODA specific activity. (f) Effect of l-DOPA concentration on the specific activity of AmDODA. Red line corresponds to the non-linear fit to the Michaelis-Menten equation. Red region indicates 95% confidence band. Standard reaction conditions: [AmDODA] = 1 μM, [AscH] = 10 mM, [DOPA] = 1 mM, sodium phosphate buffer (50 mM, pH 8.5).

#### 3.2.2. Determination of catalytic constants

The addition of l-DOPA to the biocatalytic system promptly results in the appearing of a broad absorption band with a maximum at 414 nm and shoulders at around 380 nm and 450 nm, which have been attributed, respectively, to 2,3- and 4,5-seco-DOPAs (Saha *et al*., 2015) and to betaxanthins (Fig. 3a). The enzyme activity measured via absorption spectroscopy refers to the overall conversion of l-DOPA into several products because the band centered at 414 nm may refer to betalamic acid, muscaflavin, OBDC (Saha *et al*., 2015), and any other substance showing high molar absorption coefficient at this wavelength. Although temporal changes in the absorption at 414 nm was used to investigate the catalytic activity of AmDODA and this approach has been frequently used to investigate l-DOPA dioxygenases in general (Girod & Zryd, 1991; Terradas & Wyler, 1991; Mueller *et al*., 1997b; Gandía-Herrero & García-Carmona, 2012; Contreras-Llano *et al*., 2019), one should be aware that several products and intermediates show high absorbance at this wavelength.

Attempts to separate the contribution of each species to the spectrophotometric response using second derivative spectroscopy allowed us to identify three main components of the absorption spectra (Fig. 3b) and to trace their kinetic profile (Fig. 3c), but not to discriminate between betalamic acid and muscaflavin, whose absorption spectra are superimposed. As previously reported for DODA_AMAMU (Terradas & Wyler, 1991), AmDODA can catalyze the oxidation of D-DOPA into the (4*R*)-isomer of betalamic acid, viz., isobetalamic acid (Fig. 3d).

The optimum pH for a maximum specific activity is 8.5 (Fig. 3e), which matches the conditions used for the native enzyme from *A. muscaria* (Girod & Zryd, 1991). Other l-DOPA dioxygenases, such as 4,5-DOPA-extradiol-dioxygenase of beetroots (*Beta vulgaris* L.) (Gandía-Herrero & García-Carmona, 2012), as well BmDODA1 of the silkworm (*Bombyx mori* L.) (Wang *et al*., 2019) and the YgiD protein from the bacteria *E. coli* have optimum activity at pH 8.0, whereas the 4,5-dioxygenase of *Gluconacetobacter diazotrophicus* show optimal activity at pH 6.5 (Contreras-Llano *et al*., 2019).

At pH 8.5 and 25 °C, values of *K*_M_ and V_max_ are 4.2 ± 0.4 mM and 2.6 ± 0.1 mM min^− 1^ (Fig. 3f), representing a turnover number (*k*_cat_) of 54.6 ± 0.1 min^−1^ (0.9 s^−1^) and a specificity constant (*k*_cat_/*K*_M_) of 13.0 ± 0.1 min^−1^ mM^−1^ (2.2 × 10^−4^ s^−1^ M^−1^). The value of *k*_cat_ of AmDODA in these experimental conditions is significantly lower than that of enzymes operating in the primary metabolism (*k*_cat_ ∼ 80 s^−1^) and lower than P450 enzymes (*k*_cat_ < 5 s^−1^) (Bar-Even & Salah Tawfik, 2013). The value of *K*_M_ of AmDODA is comparable to that reported for the native enzyme, i.e., 3.9 mM (Girod & Zryd, 1991) and for DODA_AMAMU, i.e. 4.5 mM (Mueller *et al*., 1997b). Nevertheless, it is lower than the values determined for the 4,5-DODA of *B. vulgaris* (BvDODA, *K*_M_ = 6.9 mM) (Gandía-Herrero & García-Carmona, 2012) and the protein YgiD from *E. coli* (*K*_M_ = 7.9 mM) (Gandía-Herrero & García-Carmona, 2013), indicating a higher affinity of AmDODA for l-DOPA compared to these dioxygenases. Furthermore, the enzyme activity in the presence of l-DOPA (0.5 U mg^−1^) is higher compared to D-DOPA (0.3 U mg^−1^). Finally, the activity of the holoenzyme and the apoenzyme were compared. Incubation of AmDODA with Chelex 100 reduces its activity over time and no catalytic conversion is observed after roughly 4 h (Fig. S6), indicating the presence of a metal cofactor that is quenched by Chelex-100. Attempts of restoring the enzyme activity by the addition of different metals, such as Fe^2+^, Fe^3+^, Mn^2+^, Zn^2+^, were not successful probably due to enzyme denaturation upon removal of the cofactor.

#### 3.2.3. Insights on the mechanism of l-DOPA oxidation

The oxidation of l-DOPA by oxygen in the presence of AmDODA and AscH was investigated using HPLC-PDA-ESI-qTOF-MS analysis. After 2 h at room temperature, and a freezing-thawing cycle, the analysis revealed four chromatographic peaks that were attributed according to the detected mass-to-charge ratio (*m*/*z*), comparison with the retention time (*R*_t_) of standard betalamic acid and l-dopaxanthin, and by literature data on muscaflavin (Mueller *et al*., 1997b; Gandía-Herrero & García-Carmona, 2013) (Fig. 4a). Although we have found muscaflavin and betalamic acid, there is no evidence on the production of OBDC (Figs. 1 and 4a). These results are corroborated by infrared multiple photon dissociation (IRMPD) analysis of the oxidation o l-DOPA in the presence of *A. muscaria* extract, which allowed the characterization of both isomers (Penna *et al*., 2020). Indeed, not all l-DOPA 2,3-dioxygenases produce all three isomers. For example, the putative l-DOPA 2,3-dioxygenases of *Streptomyces refuineus* and *Streptosporangium sibiricum* bacteria produce OBDC, but not muscaflavin or betalamic acid (Saha *et al*., 2015).

**Fig. 4.**
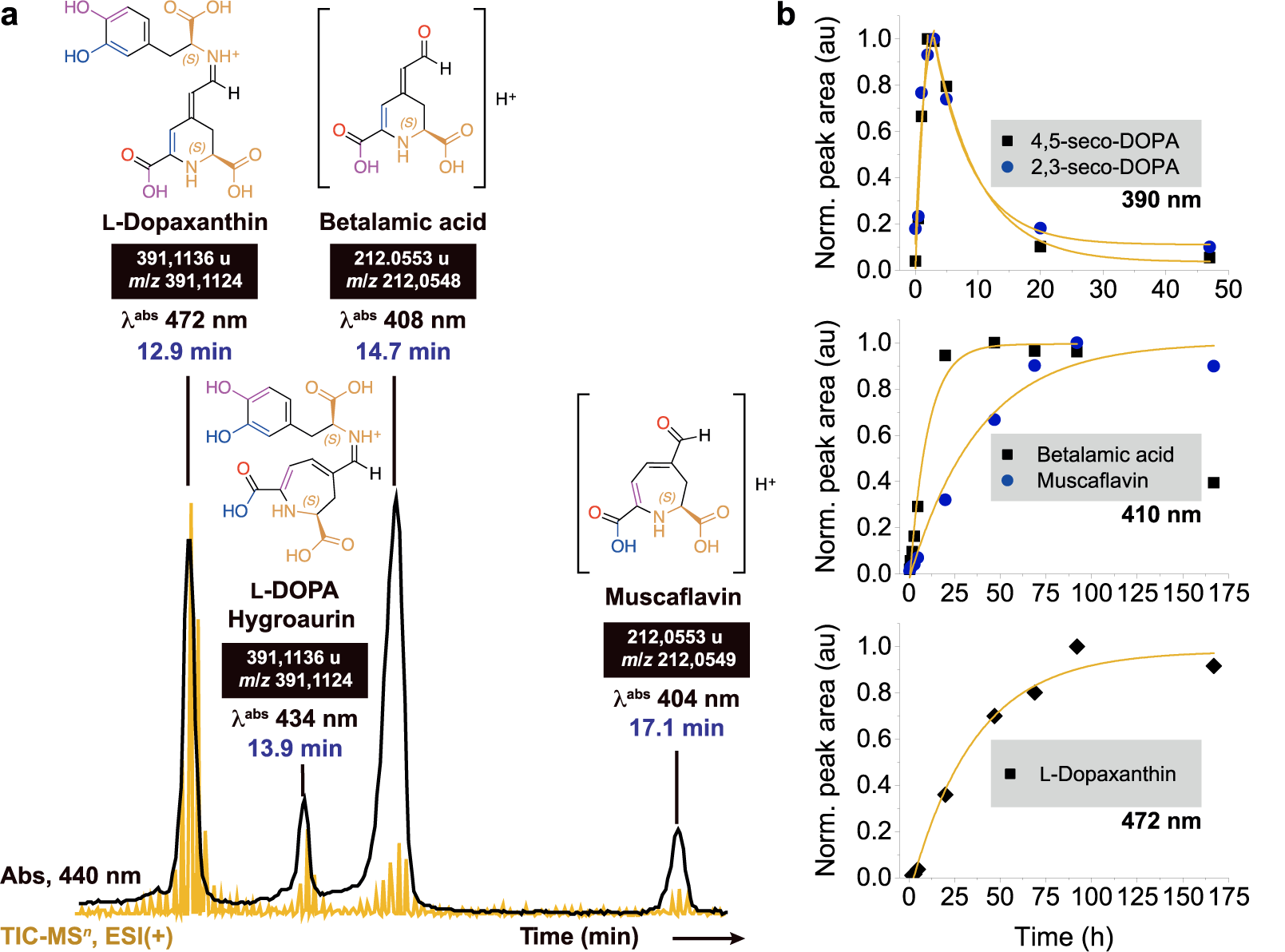
Analyses of intermediate and product formation in the oxidation of l-DOPA by oxygen in the presence of AmDODA. (a) Chromatogram of products formed after 2 h at room temperature and a freezing-thawing cycle obtained using the chromatographic condition 2.

The reaction was repeated at room temperature in the dark and monitored for 1 week at multiple times (Fig. 4b). The kinetic trace of each species was obtained by monitoring the temporal change of the area of chromatographic peaks obtained at specific wavelengths, viz., at 390 nm, 410 nm and 472 nm for the seco-DOPAs, muscaflavin/betalamic acid, and l-dopaxanthin, respectively. From these results we confirmed that the shoulder at roughly 380 nm observed in the UV-Vis experiments (Fig. 3a) corresponds to both 2,3- and 4,5-seco-DOPAs (*R*_t_ = 8.5 and 8.1 min, respectively, Fig. S7). Under the experimental condition used, these intermediates are formed for up to ca. 5 h, when their conversion into products becomes more relevant (Fig. 4b). Betalamic acid, muscaflavin and l-dopaxanthin show a sigmoid kinetic profile that agrees with the existence of precursor intermediates. The peak corresponding to the l-DOPA hygroaurin was not found under these experimental conditions, possibly because these pigments are too labile (Musso, 1979) and the sample was not submitted to a freezing-thawing cycle, which is known to promote the regeneration of hydrolyzed betalains (Herbach *et al*., 2006; Stintzing & Carle, 2007; Moreno *et al*., 2008) and could favor the coupling of muscaflavin and l-DOPA.

Raw data is presented in Fig. S4. Betalamic acid was extracted from hydrolyzed beetroot juice and purified by semi-preparative HPLC and l-dopaxanthin was semisynthesized according to the method described by Schliemann and coworkers (Schliemann *et al*., 1999), both compounds were used as HPLC standards. (b) Kinetic traces of the reaction products formed in the enzymatic oxidation of l-DOPA by oxygen in the presence of AmDODA during up to 7 d of reaction. Chromatograms, peaks retention time, absorption spectra and reaction conditions are shown in the Fig. S7 and were obtained using the chromatographic condition 1. MS^2^ spectra and ion fragments of muscaflavin and betalamic acid are presented in Fig. S8. Reaction conditions: [AmDODA] = 1 μM, [AscH] = 10 mM, [l-DOPA] = 2.5 mM, sodium phosphate buffer (50 mM, pH 8.5). The observed rate constants (*k*_obs_) obtained by fitting the data to exponential functions are presented in Table S1.

Despite the importance of l-DOPA dioxygenases for bacteria, fungi and plants and the obtaining of several recombinant examples (Contreras-Llano *et al*., 2019), details on the structure and mechanism of catalysis of these enzymes are still lacking. The concentration of the species found by HPLC-PDA-ESI-qTOF-MS analysis cannot be easily determined without knowing their molar absorptivity coefficients and/or using calibration curves for each compound. Hence, instead of using absolute concentrations, we created a simplified kinetic model based on the following constraints: *i*) 2.5 mM l-DOPA is converted into both 2,3- and 4,5-seco-DOPA, *ii*) the spontaneous and intramolecular cyclization of the seco-DOPAs produces muscaflavin and betalamic acid, *iii*) the reaction of betalamic acid with l-DOPA results in l-dopaxanthin (Kobayashi *et al*., 2001), which was originally found in the petals of *Glottiphyllum longum* (Haw.) N.E.Br. (Aizoaceae) (Impellizzeri *et al*., 1973), and *iv*) the experimentally observed rate constants (*k*_obs_) were used as first-order rate constants for each elementary reaction of the model (Table S1).

The resulting initial kinetic model could semi-quantitatively reproduce the kinetic profile observed experimentally and, most importantly, was used to convert absorption data from the chromatographic experiments into relative concentrations. The resulting rate constants are strongly dependent on the factors used to convert the area under the chromatographic peak into relative concentrations. However, this approach was useful to examine and compare parallel reactions as well as to develop a more complex kinetic model. Eleven irreversible elementary reactions were necessary to fit the kinetic data properly (*r*^2^ > 0.99, Fig. 5a). We assume that l-DOPA is in large excess over the concentration of betalamic acid and muscaflavin, and that oxygen is the limiting reagent. Also, each elementary reaction is described as irreversible to decrease the number of variables of the model. The oxidation of 2.5 mM l-DOPA by oxygen (1.0 mM) in the presence of AmDODA leads to 2,3-seco-DOPA twice as fast than to 4,5-seco-DOPA (Fig. 5b). Since the experimental values of *k*_obs_ for the consumption of 4,5-seco-DOPA does not match to that of formation of betalamic acid (3.88 × 10^−5^ s^−1^ vs. 8.58 × 10^−6^ s^−1^, respectively, Fig. 4b), we assume this transformation might involve an intermediate species. Both 2,3- and 4,5-seco-DOPAs contain a conjugated aldehyde-enol system that is prone to double keto-enol tautomerism lowering the energy barrier for conformational change. The inclusion of a tautomerization step not only explains how the geometry required for the nucleophilic attack of the amino group to the ketone carbonyl of the pyruvate moiety leading to cyclization is reached but also improves the kinetic model, as determined by the comparison of the regression coefficients. The tautomerization steps have similar rate constants for both 2,3- and 4,5-seco-DOPA. The cyclization step, however, is much faster for the conversion of 4,5-seco-DOPA into betalamic acid than the parallel reaction with 2,3-seco-DOPA, possibly because the energy of the transition state for the closure of a six-membered ring is lower than that required to produce muscaflavin, which was a seven-membered 2,3-dihydro-1*H*-azepine ring (Fig. 5b).

**Fig. 5.**
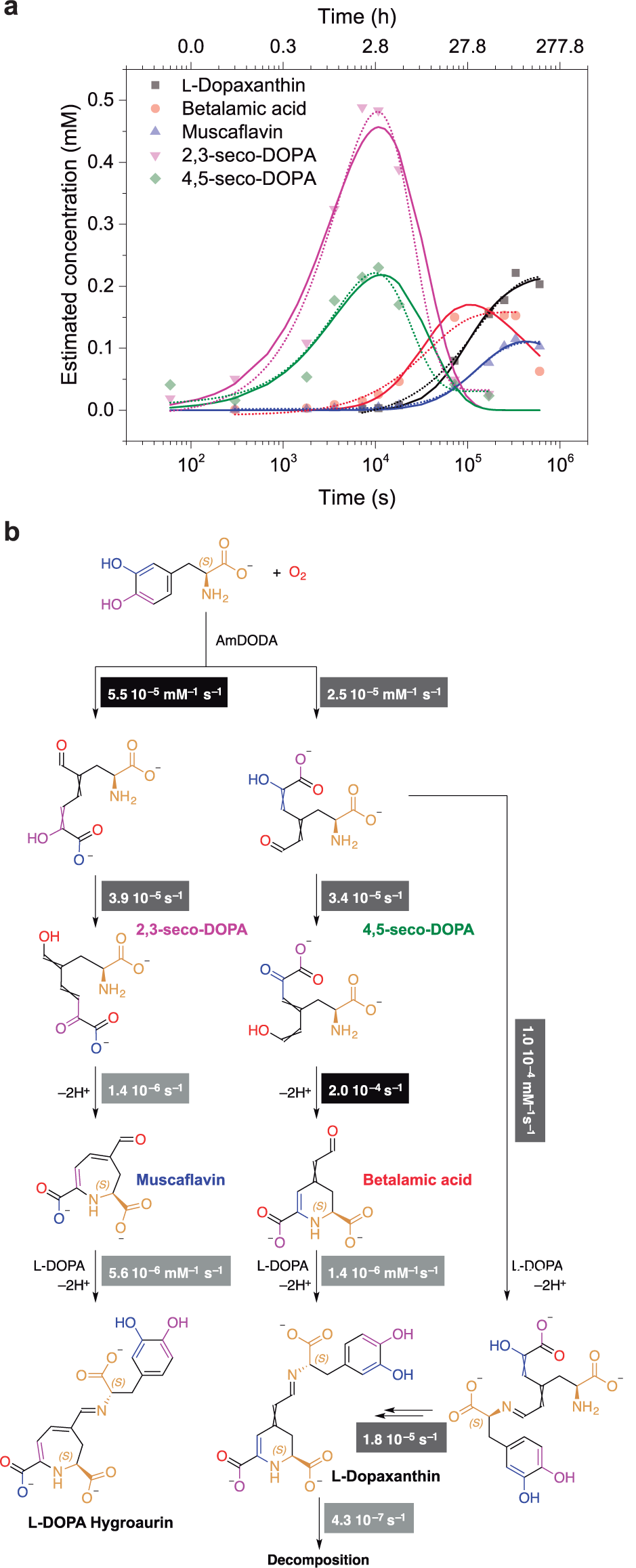
Kinetic modelling and proposed mechanism of oxidative cleavage of l-DOPA by oxygen in the presence of AmDODA.

It is important to reiterate that these rate constants are meaningful only for comparison purposes and inference of mechanistic implications. The experimental rate constant for the unimolecular cyclization of dopaquinone to cyclo-DOPA at pH 8.6, for example, is 23 s^−1^ (Thompson *et al*., 1985), which is much higher than those reported for the cyclization steps in our scheme. Furthermore, the bimolecular rate constant for the coupling of betalamic acid and cyclo-DOPA at pH 6 is < 1.7 × 10^9^ M s^−1^ (Schliemann *et al*., 1999), which for an enzyme having a *k*_cat_ ≥ 1.0 s^−1^ and operating at 50% saturation represents a reaction rate of roughly 2 × 10^−6^ M s^−1^ (Bar-Even & Salah Tawfik, 2013).

The bimolecular rate constant for the coupling between l-DOPA and betalamic acid is lower than that of the analogous reaction with muscaflavin. However, the experimental *k*_obs_ for the formation of l-dopaxanthin is not compatible with that describing the consumption of betalamic acid. The proper description of the formation of l-dopaxanthin required the addition of a parallel pathway that includes the formation of an imine adduct of 4,5-seco-DOPA and l-DOPA, followed by a cyclization step (Fig. 5b). Although the kinetics of l-DOPA oxidative cleavage by oxygen in the presence of the dioxygenases has been presented elsewhere (Terradas & Wyler, 1991), this rationalization contributes to understanding the mechanism of enzymatic catalysis allowing the comparison of AmDODA and other dioxygenases.

## CONCLUSION

We revised the CDS of the *A. muscaria*’s l-DOPA dioxygenase enabling the heterologous expression of AmDODA, which can be used to produce betalains and hygroaurins. The isomeric biosynthetic seco-DOPA precursors of betalamic acid and muscaflavin were produced by the oxidative cleavage of l-DOPA by molecular oxygen at pH 8.5 in the presence of AmDODA, which was two distinct His-His-Glu motifs in the putative catalytic pocket. The cleavage of the catechol of l-DOPA at the 2,3 position in the presence of AmDODA is faster compared to the 4,5 position. Conversely, cyclization of the 4,5-seco-DOPA leading to betalamic acid is much faster than the cyclization of the 2,3-seco-DOPA to muscaflavin. Using the protocols described here, hygroaurins can be conveniently obtained, making possible the study of this rare class of natural pigments. Last, these results will contribute to understanding the functional convergence between plant and fungi enzymes.

## ACKNOWLEDGEMENTS

We thank Mr. Enrico Florence Stevani for collecting *Amanita muscaria* for us. This research was funded by the São Paulo Research Foundation – FAPESP (ELB, 2014/14866-2 and 2019/06391-8; COM, 2015/24760-0; DMMS, 2019/12605-0; LCPG, 2018/25842-8; CVS 2017/22501-2), the Brazilian National Council for Scientific and Technological Development – CNPq (ELB, 304094/2013-7), and Coordination for the Improvement of Higher Education Personnel (CAPES, Financial code 0001).

## AUTHOR CONTRIBUTIONS

E.L.B. and C.T.H. conceived the study. C.O.M, F.A.D., E.P., and L.C.P.G. performed chromatographic analyses. L.C.E. prepared l-dopaxanthin. C.V.S. provided the *A. muscaria* fungi. L.C.P.G. and D.M.M.S. performed kinetic studies. D.M.M.S. and F.M.M.A. cloned the *dodA* gene and expressed AmDODA. C.T.H., D.M.M.S. and C.C.O. conceived the molecular biology experiments. E.L.B. carried out *in silico* studies and kinetic modelling. E.L.B., C.T.H., D.M.M.S. and L.C.P.G. wrote the paper. All authors discussed the results and revised the manuscript.

