## Supplementary material for "L-DOPA dioxygenase of the fly agaric toadstool: revision of the *dodA* gene sequence and mechanism of enzymatic pigment production"

### MATERIAL AND METHODS

#### General

All chemicals were purchased from Sigma-Aldrich and used without further purification, except as otherwise stated. Solutions were prepared using deionized water (18.2 MΩ cm at 25 ºC, TOC ≤ 4 ppb, Milli-Q, Millipore). All values are expressed as mean ± standard deviation (SD) of three completely independent replicates.

#### Buffers

**Sodium phosphate buffer** (50 mM, pH 7.4) was prepared by dissolving the appropriate amount of phosphate sodium salt in water and adjusting the pH to 7.4 with NaOH at 25 ºC. **Sodium phosphate lysis buffer** (50 mM, pH 7.4, 0.3 M NaCl) the phosphate buffer was supplemented with imidazole (10 mM), phenylmethane sulfonyl fluoride (PMSF, 1 mM), β-mercaptoethanol (2 mM) and one tablet of cOmplete™ protease inhibitor cocktail (Roche). **Elution buffer** was prepared from the sodium phosphate lysis buffer, supplemented with a linear gradient of imidazole from 100 mM to 500 mM.

**Standard compounds**

**Betalamic acid (HBt)** was obtained according to the procedure described by Schliemann and coauthors (Schliemann *et al.*, 1999) with few modifications. Fresh beetroot juice (200 mL) was paper-filtered and submitted to alkaline hydrolysis using $NH_4OH$ (pH 11.4, 30 min, room temperature (rt)). Next, the solution was cooled with an ice bath and the pH was adjusted to 3 by slow addition of concentrated HCl. Betalamic acid was extracted with ethyl acetate (50 mL, 2×) and submitted to chromatographic and spectrophotometric analysis.

**L-Dopaxanthin** was obtained as described by Schlieman and coauthors (Schliemann *et al.*, 1999), with few modifications. Betalamic acid in ethyl acetate was obtained from hydrolyzed beetroot juice. The acid was partitioned into water and the residual organic solvent was evaporated under reduced pressure (80 mmHg, 25 ºC). L-DOPA (50 equiv.) was added to the solution of HBt (5 mL) in water pH 10 and the depletion of HBt and concomitant appearance of the L-dopaxanthin were monitored spectrophotometrically at 430 nm and 470 nm, respectively. After completion, HCl (*conc.*) was added slowly until pH 5 was reached. The product was purified by gel chromatography using Sephadex LH-20 as stationary phase and water as eluent. Final characterization was carried out by LC-HRMS.

### FIGURES

```
   1 acatgcgctt cgtcactcaa atatatccat tggtttcgcg acacgctggt tccgaaccca
  61 gagagtctcg cgactctctt cagaaattct cccaacttgc gttccttcaa cacccctttg
 121 gtgcccgcgg cagaggactg cgacagttag ccatgacgga ttctttcatt cctctctgac
 181 gactatccat gcagaggcca ggattctgga ctatgctagt cctatcccac ctcatctcat
 241 gaaccttcct tctctacgcc agatgtcctt cgttgttcct ctttcgccgg gcagacgcac
 301 gttggcgcag tttcctcgag tggcatggtt cacgtttaca gcatatccaa gctttgattt
 361 accttgcagg gaaccacttc cagtccgact tggcctacat tgcatcatca tgccgcaacc
 421 ttcttagact cgatcttgct ttctcccatt ggacggcata tgccgtttcc tgttagcctc
 481 ccgcctacag tggaacatct gggcatcttt tgtactcagg gtcagatcgc caactaccag
 541 acgttcttct ctcgtttaga taggatcgag tatggtgcca agcttcgttg tatacagttc
 601 ttgggtaaac ggacgtcaga gatatattcg acaagcattt gcatcaattc tggtctggca
 661 cgaggcgact aagaggtagt ggggttagat tcatgaatca tagaggggca tgtatagcgt
 721 agctaaggat agcactggca gttccacagt tctatataat ccgtgatacg actctctcgt
 781 tcccctcaca tactaccatg tccaccaagc cagagactga ccttcaaact gtcctcgaca
 841 gcgaaatcaa gtttgtctag tttgtctcca ctagcaaaaa ctctgctgac ttgtgtacag
 901 ggaatggcac tttcgtcagt cgaccaagca acgactaact tatgtagctc aaattgaccc
 961 tgcaactaat agacatctac tttcatcaga acaacgccgc agagcatcaa gctgcgcttg
1021 agcttcgtga cgcggttctg aggctcagac aagacggcgc attcgtcgcc gttcccttgt
1081 tccgcgttaa catggacccc atgggtcctc atcctgtcgg tcagtagtca tcgcaatcac
1141 accaccagtc cttgaacaag tcccttgact ctcaaggttc ttatgagatc tgggttccgt
1201 ctgaaacgtt cgcttccgtg ttctcctact tgtgcatgaa cagagggaga ttaagcatcc
1261 ttgtgcatcc tttgacacgc gaagaactca gagaccatga aattcgtaat gcctggatag
1321 gaccctcttt cccactcaat ctcgccaacc taccgatcaa gagtgatgag atccctttgc
1381 aatatccaag cctcagtatg tcatctcacc actctctggc ggccctcagt gacaaattgt
1441 tgcagagctt gggtactcat cgacagcgca taagatgtca ttggaagaaa ggcggaaatt
1501 aggcgacgat atagaagcag tgcttagggg agagaaagag gcggccagag cgccccatcg
1561 agatgcatag agctacattc gattgtctat attgctactg acatagggta atgttggaat
1621 tttctgccc
```

**Fig. S1.** DNA sequence for *dodA* gene (Hinz *et al.*, 1997; GenBank accession Y12886, NCBI). *dodA* exons are painted with blue color and 3'-AG canonical splice site, altered to GA in AmDODA sequence, is highlighted in yellow. The start codon for AmDODA CDS is shown in red.

**M**VPSFVVYSSWVNGRQRYIRQAFASILFYIIRDTTLSFPSHTT**M**STKPETDLQTVLDSEIKEWHFHIYFHQNN

AAEHQAALELRDAVLRLRQDGAFVAVPLFRVNMDPMGPHPVGSYEIWVPSETFASVFSYLCMNRGRLSILVHP

LTREELRDHEIRNAWIGPSFPLNLANLPIKSDEIPLQYPSLKLGYSSTAHKMSLEERRKLGDDIEAVLRGEKE

AARAPHRDA-

**Fig. S2.** Amino acid sequence for DODA_AMAMU (full sequence) and AmDODA (sequence in yellow).

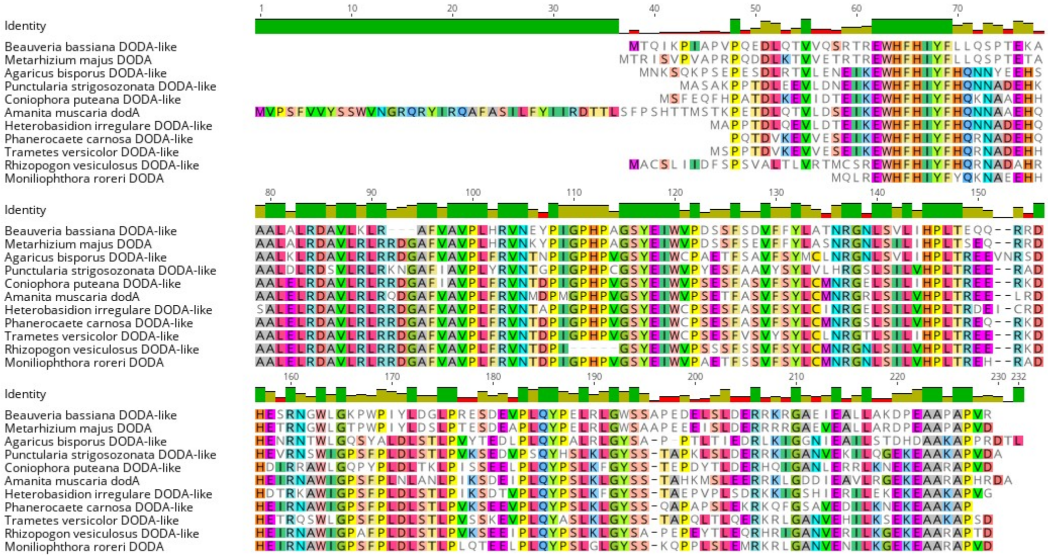

**Fig. S3.** Multiple alignment of amino acid sequences of fungal L-DOPA-dioxygenases (MUSCLE, Geneious Prime® v11.0.4).

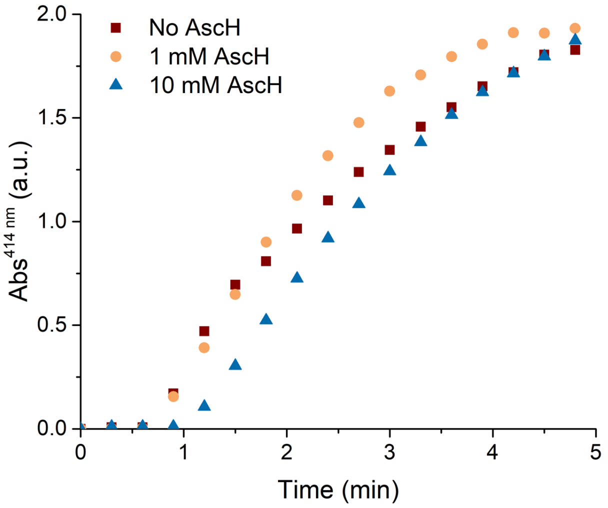


**Fig. S4.** Effect of ascorbic acid (AscH) on the oxidative cleavage of L-DOPA by oxygen catalyzed by AmDODA. Absorption at 414 nm of the conversion of L-DOPA (1 mM) by AmDODA (1 µM) in the presence and absence of 1 or 10 mmol $L^{-1}$ AscH in sodium phosphate buffer (50 mM), pH 8.5.

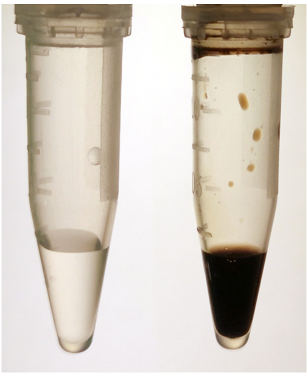

**Fig. S5.** Effect of the presence (left) and absence (right) of ascorbic acid (AscH) in the standard reaction without AmDODA.

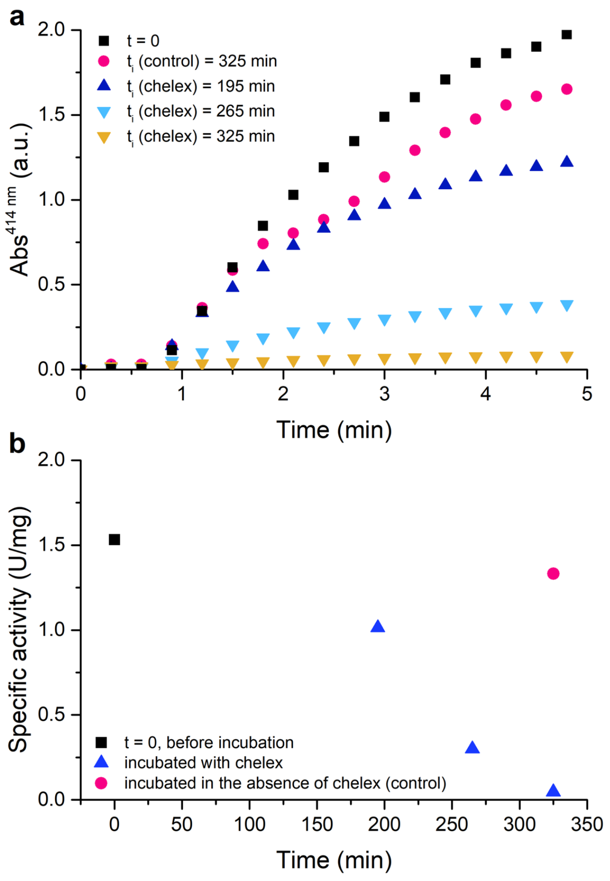


**Fig. S6.** Effect of the incubation with Chelex-100 on the oxidative cleavage of L-DOPA by oxygen catalyzed by AmDODA. (a) Absorption at 414 nm of AmDODA (0.24 mg/mL) incubated at 4 °C under agitation (450 rpm), with and without Chelex-100 in sodium phosphate buffer (50 mM), pH 8.5. (b) Specific activity (U/mg) at the same conditions. Reaction condition: [AmDODA] = 1 μM, [AscH] = 10 mM, [L-DOPA] = 1 mM, sodium phosphate buffer (50 mM), pH 8.5.

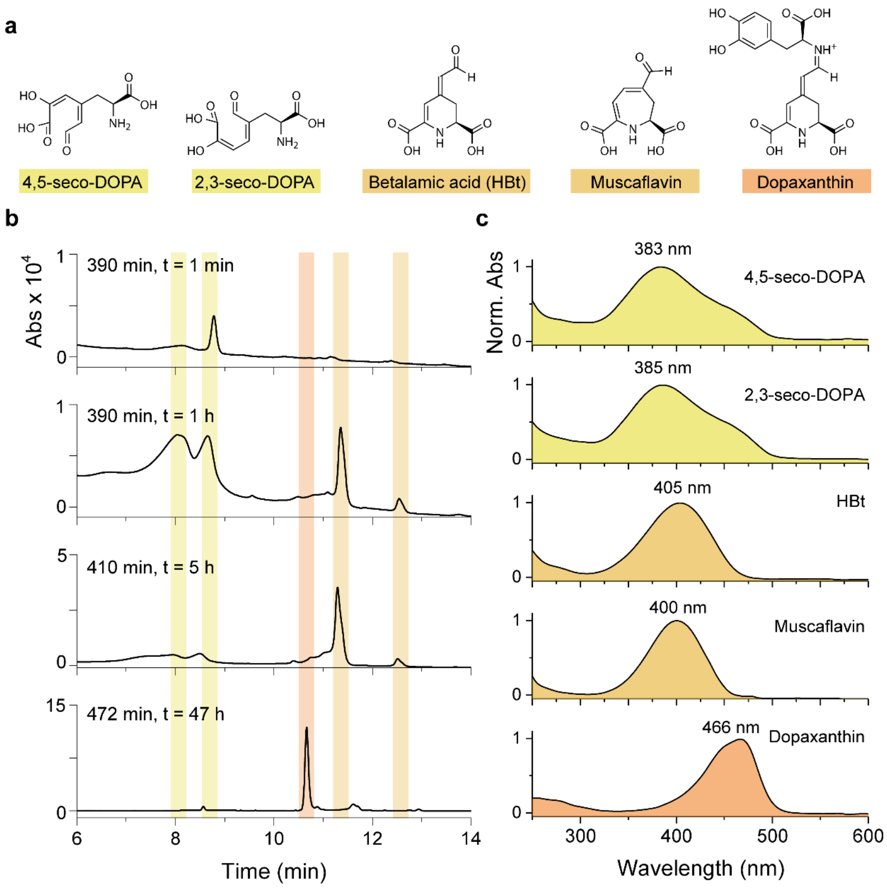


**Fig. S7.** HPLC analysis of 4,5-seco-DOPA and 2,3-seco-DOPA formed during oxidative cleavage of L-DOPA by oxygen catalyzed by AmDODA and derived compounds (betalamic acid, muscaflavin and dopaxanthin). (a) Reaction products monitored by HPLC analysis. (b) Chromatograms obtained at 390 nm (seco-DOPA), 410 nm (betalamic acid and muscaflavin) and 472 nm (dopaxanthin). Note that *t* corresponds to the time of injection after the reaction was triggered. (c) Spectra of the peaks of the reaction products with retention times: 8.1 min (4,5-seco-DOPA), 8.7 min (2,3-seco-DOPA), 10.7 min (L-dopaxanthin), 11.3 min (betalamic acid), 12.5 min (muscaflavin).

Reaction condition: [AmDODA] = 1 µM, [AscH] = 10 mM, [L-DOPA] = 2.5 mM, sodium phosphate buffer (50 mM), pH 8.5.

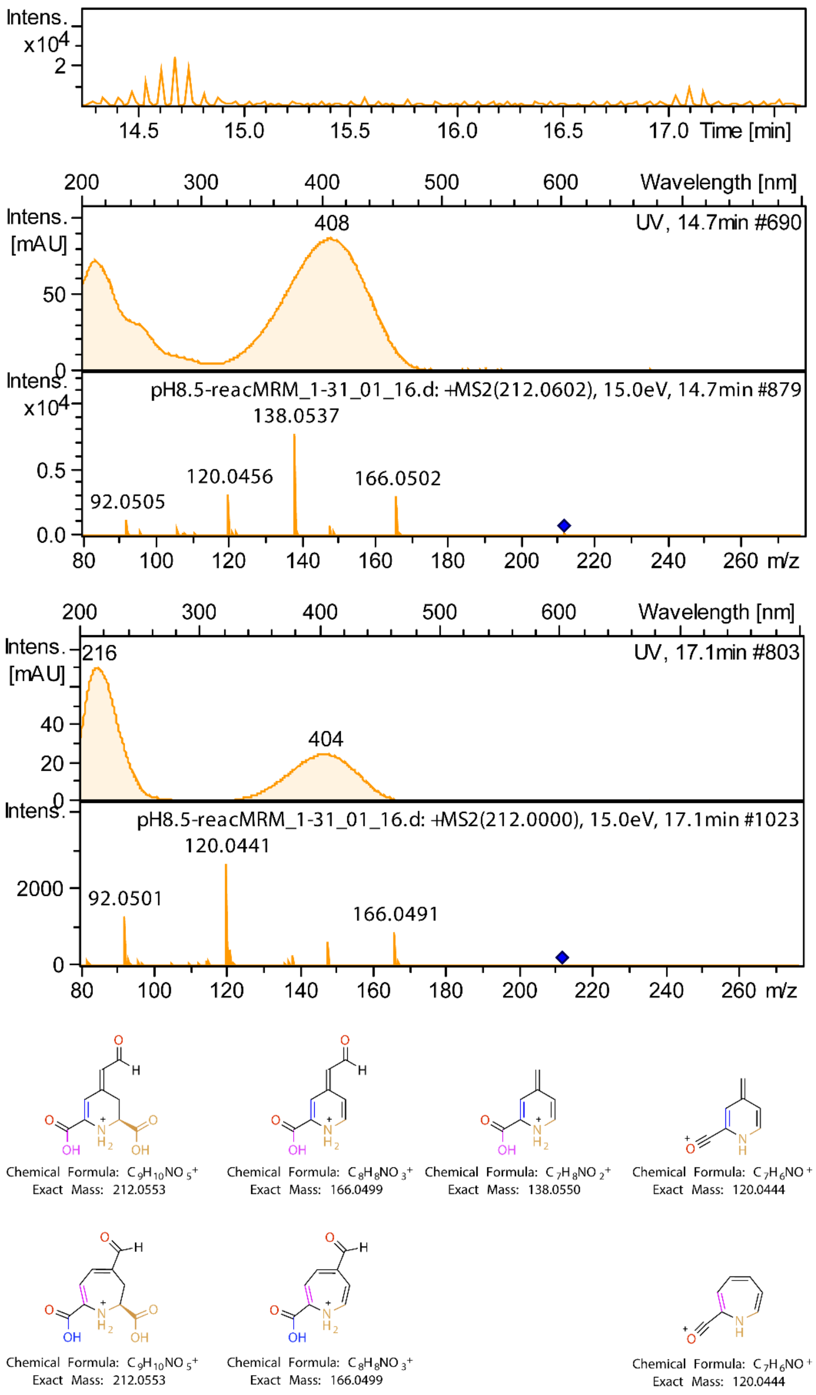


**Fig. S8.** Comparison of HPLC-PDA-MS(ESI) and MS/MS analysis of the substances with retention time of 14.7 min and 17.1 min, which were characterized as betalamic acid and muscaflavin.

### TABLE

**Table S1.** The observed rate constants ($k_{obs}$) calculate by fitting the data (normalized peak area *vs.* time) to exponential functions. The $k_{obs1}$ refers to the exponential decay and $k_{obs2}$ to the exponential growth.

| **Compound** | **$k_{obs1}$ ($s^{-1}$)** | **$k_{obs2}$ ($s^{-1}$)** |
|---|---|---|
| 4,5-seco-DOPA | $3.9 \times 10^{-5} \pm 8.9 \times 10^{-8}$ | $9.1 \times 10^{-5} \pm 4.5 \times 10^{-7}$ |
| 2,3-seco-DOPA | $4.5 \times 10^{-5} \pm 1.2 \times 10^{-7}$ | $2.0 \times 10^{-4} \pm 1.9\times 10^{-6}$ |
| Betalamic acid | | $2.8 \times 10^{-5} \pm 4.6 \times 10^{-8}$ |
| Muscaflavin | | $7.3 \times 10^{-6} \pm 3.1 \times 10^{-9}$ |
| L-Dopaxanthin | | $7.9 \times 10^{-6} \pm 3.6 \times 10^{-9}$ |
